# Efficient parameterization of large-scale mechanistic models enables drug response prediction for cancer cell lines

**DOI:** 10.1101/174094

**Authors:** Fabian Fröhlich, Thomas Kessler, Daniel Weindl, Alexey Shadrin, Leonard Schmiester, Hendrik Hache, Artur Muradyan, Moritz Schütte, Ji-Hyun Lim, Matthias Heinig, Fabian J. Theis, Hans Lehrach, Christoph Wierling, Bodo Lange, Jan Hasenauer

## Abstract

The response of cancer cells to drugs is determined by various factors, including the cells’ mutations and gene expression levels. These factors can be assessed using next-generation sequencing. Their integration with vast prior knowledge on signaling pathways is, however, limited by the availability of mathematical models and scalable computational methods. Here, we present a computational framework for the parameterization of large-scale mechanistic models and its application to the prediction of drug response of cancer cell lines from exome and transcriptome sequencing data. With this framework, we parameterized a mechanistic model describing major cancer-associated signaling pathways (>1200 species and >2600 reactions) using drug response data. For the parameterized mechanistic model, we found a prediction accuracy, which exceeds that of the considered statistical approaches. Our results demonstrate for the first time the massive integration of heterogeneous datasets using large-scale mechanistic models, and how these models facilitate individualized predictions of drug response. We anticipate our parameterized model to be a starting point for the development of more comprehensive, curated models of signaling pathways, accounting for additional pathways and drugs.

## Introduction

Personalized tumor therapy relies on our ability to predict the drug response of cancer cells from genomic data^1^. This requires the integration of genomic data with available prior knowledge, and its interpretation in the context of cancer-associated processes^2^. At the heart of this endeavor are statistical and mechanistic mathematical models^3^. In patient stratification, statistical models are used to derive prognostic and predictive signatures of tumor subtypes^4^,^5^. Linear and nonlinear regression, machine learning methods and related approaches have been used to obtain such signatures^6^. Yet, purely statistical models do not provide mechanistic insights or information about actionable targets. High-quality mechanistic models of cancer signaling are thus of interest to researchers and clinicians in systems biology and systems medicine.

Mechanistic models aim to quantitatively describe biological processes. Consequently, they facilitate the systematic integration of prior knowledge on signaling pathways, as well as the effect of somatic mutations and gene expression. These models have been used for the identification of drug targets^7^ as well as the development of prognostic signatures^8^,^9^. Furthermore, mechanistic modeling has facilitated the study of oncogene addiction^10^, synthetic-lethal phenotypes^11^ and many other relevant phenomena^12^.

Various pathways have been modeled mechanistically using Ordinary Differential Equations (ODEs) of varying detail^13^. ERBB, MAPK and PI3K signaling attracted special attention as they are altered in many cancer types^14^ and targeted by many drugs^15^. Tailoring the models to individual pathways ensures manageability of the development effort, but neglects crosstalk. The Atlas of Cancer Signaling Network (ACSN) addresses this issue by covering a majority of molecular processes implicated in cancer^16^. However, like other pathway maps^17,18^, the ACSN lacks kinetic rate laws and rate constants, preventing numerical simulation and quantitative prediction. This might be addressed in the future by using comprehensive databases^13,19,20^ in combination with semi-automatic^21–23^ or automatic reconstruction methods^8,24,25^.

After the construction of a mechanistic model, parameterization from experimental data is necessary to render the model predictive. Optimization methods achieve this by iteratively minimizing the objective function, i.e. the distance between model simulation and experimental data^26,27^. This requires repeated numerical simulations. As even for medium-scale models millions of simulations are necessary, the computational burden is often immense^28^. Accordingly, parameterizing large-scale pathway models is often deemed intractable and has not been done in practice. A scalable method for parameterization of large-scale mechanistic models is therefore essential for the community as it enables the comprehensive integration of prior knowledge and experimental data.

Here, we describe a large-scale mechanistic model of cancer signaling which is individualized using information about somatic mutations and gene expression levels. We introduce a computational framework for the parameterization of large-scale ODE models that reduces the computation time by multiple orders of magnitude compared to state-of-the-art methods. We demonstrate the parameterization of the model from thousands of drug assays from over 100 human cancer cell lines and validate the predictive power of the model. Moreover, we show that the mechanistic model outperforms all investigated statistical models in terms of classification accuracy and generalizes to cancer cell lines originating from tissues not used for training.

## Results

### Large-scale mechanistic model integrates knowledge of cancer signaling pathways

To predict the drug response of cancer cell lines, we developed a mechanistic model integrating signaling modules reflecting the human ERBB, RAS and PI3K/AKT signaling pathways, as well as regulation of the transcription factors MYC and AP1^29^ (Fig. 1a). This model describes synthesis, degradation, translocation, complex formation, phosphorylation and various other types of reactions for proteins and their functional variants (Supplementary Fig. 1). We assembled it manually using the web-based platform PyBioS30,31 and provide it as annotated Systems Biology Markup Language (SBML) file (Supplementary File 1). The model is based on curated information from ConsensusPathDB^32^, a meta-database integrating more than 20 public databases (e.g., DrugBank^33^, KEGG^34^ and Reactome^35^), and additional publications.

**Figure 1.**
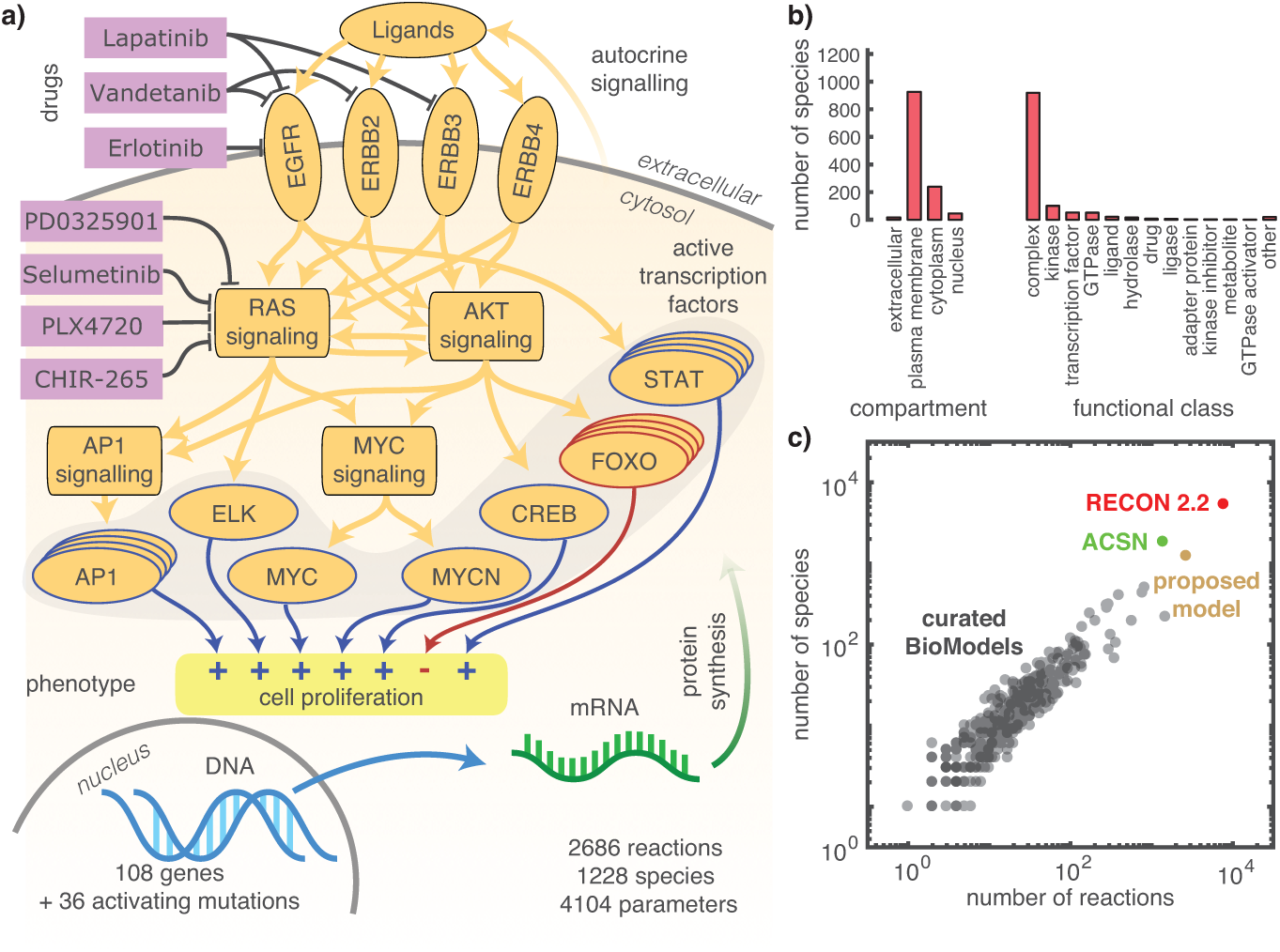
Model structure and properties. (a) Sketch of modeled signaling pathways. The developed model describes the synthesis and protein-protein interactions for protein products of 108 genes and 36 activating mutations. The visualization depicts drugs (purple), selected molecular species (orange) and cell proliferation as phenotypic readout (yellow). (b) Distribution of modeled molecular species on compartments and functional classes (c) Comparison of complexity of the proposed model with curated models from the BioModels database^13^, Recon 2.2^36^ and the ACSN^16^.

The model accounts for 108 genes and 36 activating mutations yielding a total of 1228 molecular species in 4 cellular compartments (Fig. 1b) involved in 2686 reactions. The modeled mutations cover 7 of the 10 most frequent driver mutations reported by Rubio-Perez et al.^36^ and account for 22.1% of driver mutations observed in patient samples^36^. The model describes the action of 7 different small molecule kinase inhibitors, of which 4 are FDA-approved. For 17 additional FDA-approved kinase inhibitors, one or more main targets are included in the model, but the action is currently not described. Thus, the model covers main targets for 27.3% of FDA-approved targeted cancer therapies.

To quantify the scale of our model, we compared it to curated models available in the BioModels database^13^. The proposed model describes more biochemical species and reactions than any other of the curated models (Fig. 1c). Two pathway maps, Recon 2.2^18^ and the ACSN, possess a similar size than our model. Yet these pathway maps do not provide kinetic rate laws and Recon 2.2 does not focus on cancer signaling. Hence, we conclude that the proposed model is one of the most extensive mechanistic models of cancer signaling.

Most drug response assays provide information about the cell proliferation rate relative to the untreated condition. Cell proliferation of cancer cells is governed by a complex interplay of cellular signaling processes regulating e.g. the balance between pro-growth and (anti-) apoptotic signals in response to extracellular stimuli or presence of activating mutations within respective signal transduction cascades. A major function in regulation of cell proliferation has been attributed to transcription factor (TF) activation, e.g. of the MYC, AP1 and FOXO transcription factors, and regulation of target gene expression in response to extracellular or oncogenic stimuli. In the current model we used the weighted sums of the simulated molecularly activated state of these TFs as a surrogate for proliferation (see Online Methods, Section Model Development). This semi-mechanistic description provides a simple model of down-stream regulatory processes.

### Genomic data provides basis for individualization of the mechanistic model

The mechanistic model provides a generic template for a subset of signaling processes in human cells. To obtain a model for a particular cancer cell line, we individualized the mechanistic model by incorporating gene expression levels as synthesis rates for proteins and their mutated functional variants (Fig. 2a). We assumed that all other kinetic parameters such as transport, binding and phosphorylation rates, only depend on the chemical properties of the involved biochemical species. Accordingly, these parameters differ between proteins and their functional variants, while they are assumed to be identical across cell lines. This enables the simultaneous consideration of multiple cell lines and drugs for the model parameterization, increasing the available training sample size. Furthermore, the assumption allows us to predict the drug response of new cell lines from information about gene expression levels and functional variants.

**Figure 2.**
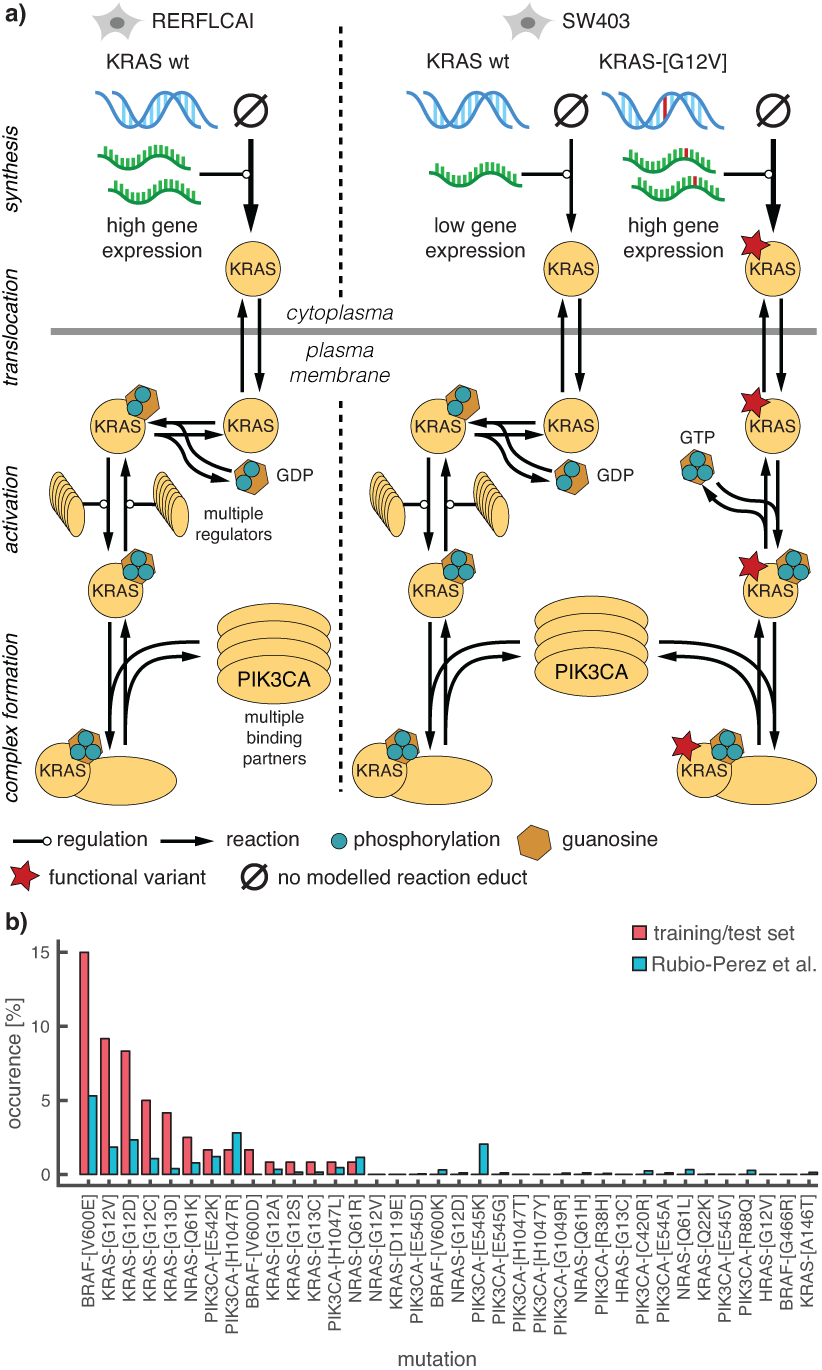
Individualization of the model with genomic and transcriptomic data. (a) Individualization of the generic mechanistic model for the two different cell lines: RERFLCAI (wild-type KRAS); and SW403 (wild-type and mutated KRAS). KRAS signaling model is illustrated from synthesis until complex formation. Degradation reactions are omitted. (b) Comparison of the occurrence frequency of mutations included in the model between the training/test set extracted from the Cancer Cell Line Encyclopedia and the InTOGen database by Rubio-Perez *et al*.^36^, which provides an extensive characterization of somatic mutations in human tumors.

In this study, we considered data for 120 human cancer cell lines from 5 tissues (breast, large-intestine, lung, pancreas and skin) provided in the Cancer Cell Line Encyclopedia (CCLE)^37^. We processed the included genetic characterization of cell lines in the untreated condition using a standardized bioinformatics pipeline (Online Methods, Section Data Processing). Of the modeled driver-mutations, 14 are present in more than one cell line (Fig. 2b).

### Scalable, parallel optimization method enables model parameterization

The mechanistic model includes more than 4,100 unknown parameters, i.e. kinetic constants and weighting factors. To describe the available data and to predict future experiments, we parameterized the model using measured proliferation data from 120 cell lines treated with 7 different drugs at up to 9 concentrations provided in the CCLE. In total, this dataset provides more than 6,900 experimental conditions. To assess the prediction uncertainty, we performed 5-fold cross-validation with 5 pairs of training (80%; 96 cell lines) and test datasets (20%; 24 cell lines).

To parameterize the model from the training data, we minimized the sum of squared residuals of measured and simulated relative proliferation. This non-linear and non-convex ODE-constrained optimization problem was solved using multi-start local optimization, an efficient and reliable approach that outperformed global optimization methods in several studies^26,38^. As the optimization problem is high dimensional, we first assessed the applicability of state-of-the-art methods, such as forward sensitivity analysis^26^. Therefore, we determined the computation time per gradient evaluation. This revealed that due to (i) the large-scale ODE model, (ii) the large number of parameters and (iii) the large number of experimental conditions, a single evaluation of the objective function gradient would require more than 5.10^4^ CPU hours (> 6 CPU years) (Fig. 3a). As the gradient has to be evaluated hundreds of times for a single optimization, available toolboxes were not applicable.

**Figure 3.**
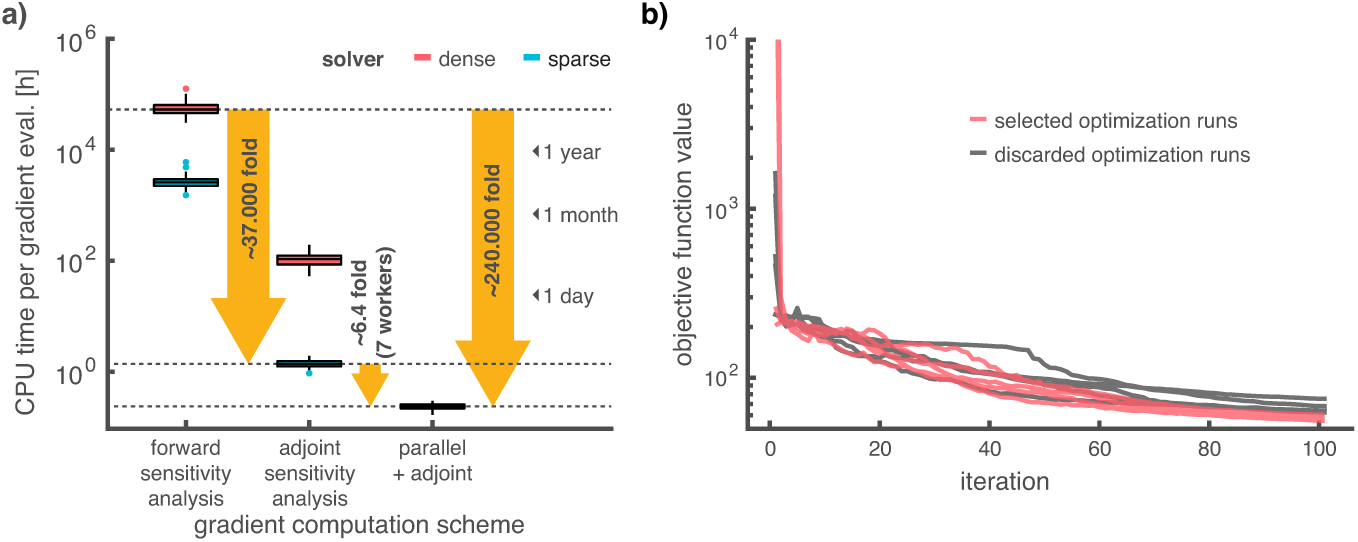
Parameterization of the mechanistic model. (a) Computation time for one evaluation of the objective function gradient, which determines the per-iteration time for a single local optimization step. For the non-parallelized evaluation, the time was computed based on representative samples (See Online Methods, Numerical Benchmark). The gradient evaluation time was dramatically reduced by using adjoint sensitivities, exploiting sparsity and parallelization. (b) Objective function traces for ten different local optimization runs for the first cross-validation set. Initial conditions for the local optimization runs are sampled from a latin hypercube. Although higher initial objective function values were observed, the corresponding axis was cropped at 10^4^. The 5 best optimization runs are colored in red and used for subsequent analysis.

To render parameterization tractable, we addressed challenges (i)-(iii). Firstly, we reduced the CPU time per model evaluation by using a sparse linear solver^39^ for ODE integration (0.5% non-zero entries in the Jacobian). Secondly, we implemented adjoint sensitivity analysis^40^ which improves scaling with the number of parameters. These two methodological advancements reduced the computation time over 37,000-fold (Fig. 3a).

Thirdly, we established scalability with respect to the number of experimental conditions by parallelization on the level of cell lines (Fig. 3a). Using 8 cores (7 workers), we observed a 6.4-fold acceleration. In total, our flexible and easily extendable parameterization framework reduced the expected wall time by over 240,000-fold.

Using 400 cores and a trivial parallelization over local optimizations, our computational framework enabled the parameterization for all cross-validations in less than one week. In comparison, state-of-the-art approaches would have required hundreds of thousands of years. The local optimization achieved a substantial reduction of the sum of squared residuals within a few iterations and then the curve flattened out (Fig. 3b). The optimization was stopped early at 100 iterations to improve the prediction accuracy by avoiding overfitting^41^. To filter insufficient optimization runs and improve robustness, we used ensemble averaging (see Online Methods, Section Ensemble Averaging) over the 5 optimization runs that achieved lowest objective function value in each cross-validation for all following analysis and prediction.

### Mechanistic model yields quantitative description of experimental data and generalizes to test data

The parameterized model describes the drug dose-dependent proliferation of cell lines. To assess the combined quality of the model and the parameterization, we quantified the model-data mismatch. Our study of the parameterized model revealed a good agreement of measured data and model simulation and little variation in the prediction (Fig. 4a), despite large parameter uncertainties (Supplementary Fig. 2). For the highest drug concentration (8µM) of each dose response curve, the correlation of measured and simulated proliferation was r=0.82 (p<10^-150^) (Fig. 4b, for more concentrations see Supplementary Fig. 3).

**Figure 4.**
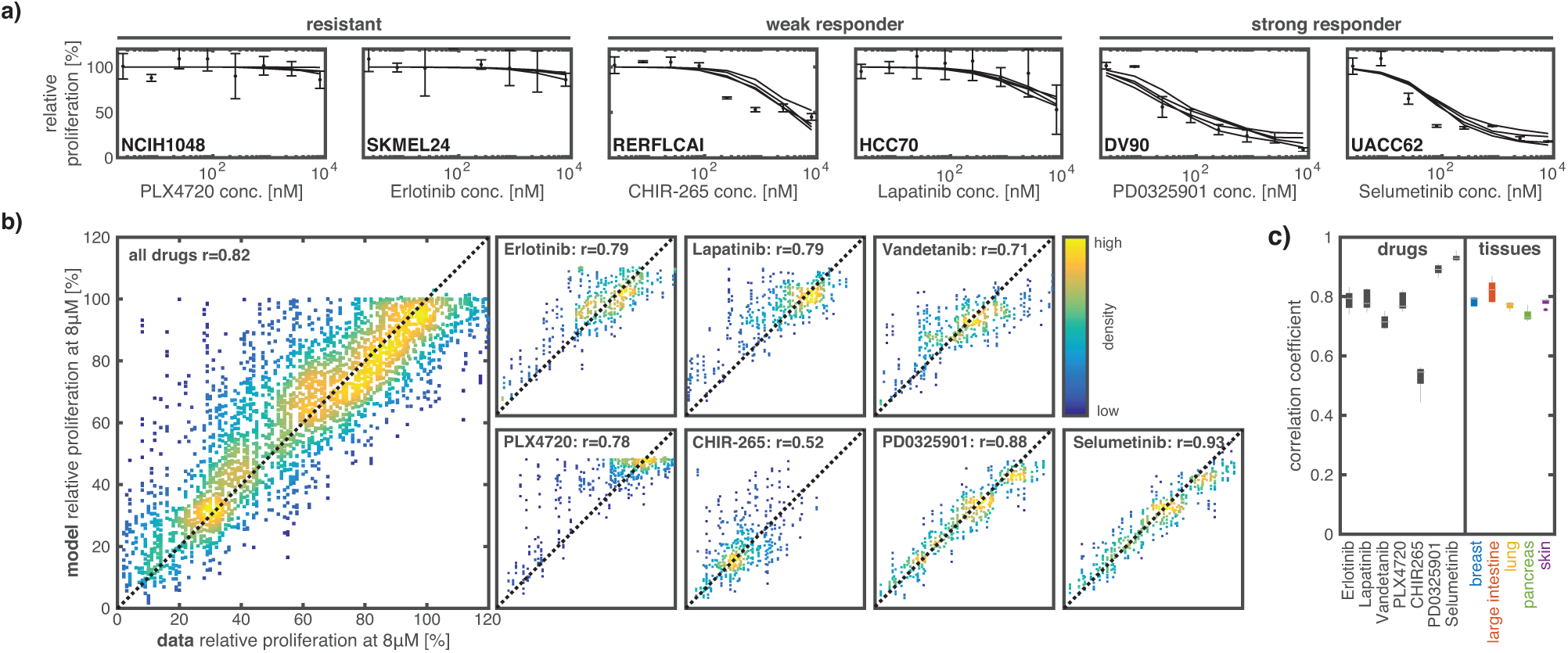
Analysis of fitting properties of the model. (a) Representative examples of model simulations for six combinations of drugs (x-label) and cell lines (bold text on bottom, left). The five plotted lines are the median fit for the five best optimization runs for every cross-validation set. (b) Correlation of simulation and measured proliferation for the response at maximal concentration. For the simulation the ensemble prediction based on the median is shown. Smaller subplots show the correlation filtered for individual drugs. (c) Correlation statistics over cross-validations for individual drugs and tissues.

The agreement of measured and simulated proliferation is similar for all tissues but varies between drugs (Fig. 4c). Similarly, no dependence of the model-data mismatch on mutation status could be identified (Supplementary Fig. 4). Particularly good correlations were achieved for selumetinib (r=0.93) and PD0325901 (r=0.88). Interestingly, CHIR-265 (r=0.52) and PLX4720 (r=0.78) have distinct correlation coefficients although they share B-RAF^V600E^ as main target with similar affinity. Still, many cell lines respond to CHIR-265, but appear to be resistant against PLX4720. This suggests that the molecular understanding of the drug/target effects or the implementation in the model may be incomplete. For example, inhibition of VEGFR2 activation by CHIR-265 has been described^42^, but is not captured by the current version of the model.

To evaluate the predictive power of the parameterized mechanistic model, we turned to the test sets of the cross-validation (see Fig. 5a left). We quantified (i) the correlation of measured and predicted proliferation as well as (ii) the classification accuracy for responder cell lines in terms of the area under the receiver-operating-characteristic (ROC) and the precision-recall (PR) curve. A cell line was considered a responder to a drug when the proliferation at the highest drug concentration was below 50% compared to the untreated control. Our analysis of the correlation revealed a good quantitative agreement of measured and predicted relative proliferation for the test set (r=0.55, p<10^-8^) (Fig. 5b), a lower correlation comparing to the training set data (r=0.82, p<10^-150^) (Fig. 5b). For the qualitative predictions, we found an average classification accuracy of 76.7±1.8% (area under ROC=0.767±0.018, area under PR=0.73±0.022) (Fig. 5c and Supplementary Figure 5).

**Figure 5.**
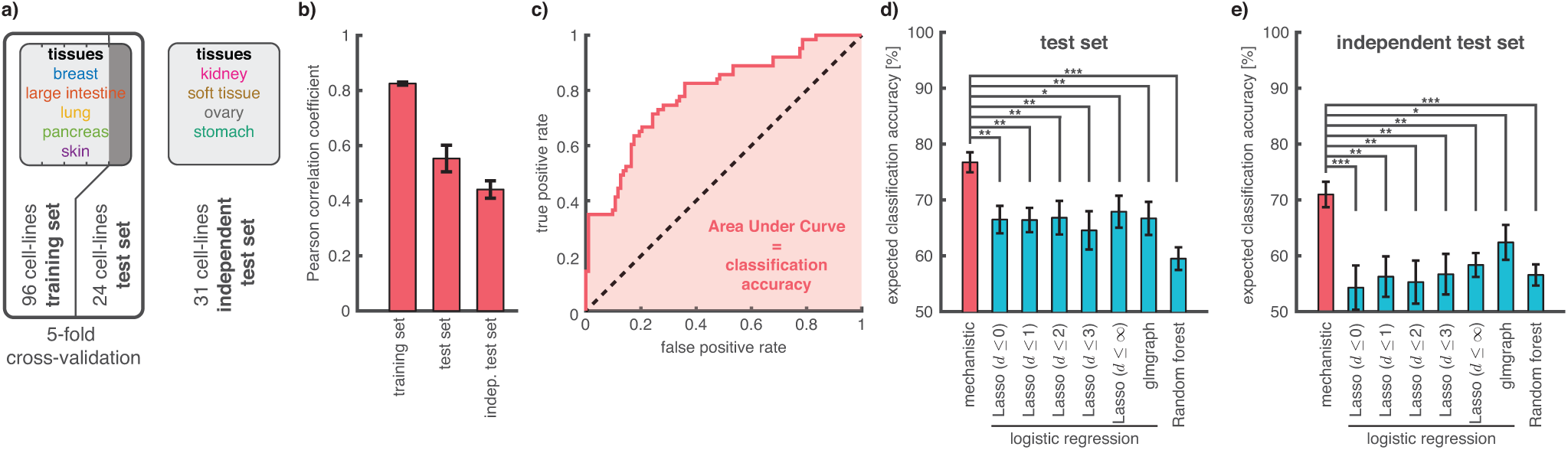
Validation of model prediction for single-drug treatment. (a) Overview over CCLE datasets used to evaluate the classification accuracy: (i) test set for cell lines originating from the same tissues on which the model was trained; and (ii) independent test set using cell lines from different tissues. (b) Correlation coefficients between measured and predicted proliferation for 8µM drug concentration on all considered datasets. Error bars show the standard error (c) Representative receiver-operating-characteristic curve for the mechanistic model on the test set (d-e) Comparison of classification accuracy across all tested methods for test set and independent test set. For the Lasso approach the d defines up to which network distance interactions are considered. Error bars show the standard error. Stars indicate statistical significance (*: p<0.05; **: p<0.01;***: p<0.001).

### Mechanistic model outperforms established statistical models

To provide a reference for the performance of the parameterized mechanistic model, we trained several well-established statistical models on the training set. The statistical models include a random forest^43^, sparse linear and nonlinear regression models^44^, as well as a network-constrained sparse regression model (with network derived from the mechanistic model)^45^. The training of all statistical models was performed using state-of-the-art toolboxes (see Online Methods, Section Statistical Analysis). The best statistical model achieved a classification accuracy of 67.8±2.9% on the test set (Fig. 5d), which is 8.9 percentage points lower than the classification accuracy of the mechanistic model (76.7±1.8%). In conclusion, the parameterized mechanistic model provides significantly (p<3.6.10^-2^ according to Welch’s *t*-test) more accurate classification than all considered statistical models.

Following these positive results, we assessed the generalization error of the mechanistic model. We processed experimental data for 31 additional cell lines from 4 additional tissues (kidney, ovary, soft tissue and stomach) available in the CCLE database that were not part of the initial training set (see Fig. 5a right). For this independent dataset, the parameterized mechanistic model achieved a classification accuracy of 70.7±2.3% (Fig. 5e), which is 6 percentage points lower than on the test set. Interestingly, the tested statistical models achieve a maximal classification accuracy of 62.1±3.1%, suggesting that our proposed parameterization framework for mechanistic models may achieve better generalization properties.

### Combination treatment outcomes predicted from single treatment measurements

For an additional assessment, we predicted the outcome of combination treatments. We considered the dataset published by Friedman *et al.*^46^ reporting the response of cancer cell lines to individual drugs as well as to combinations of two drugs. The dataset includes 7 cell lines and 5 drugs (selumetinib, CHIR-265, erlotinib, lapatinib, PLX4720) contained in our training set. To establish a reference for the accuracy of the prediction, we assessed the agreement of the proliferation measurements by Friedman *et al.*^46^ and the measurements from the CCLE database. The comparison yielded a correlation of r=0.33. This weak correlation is likely due to differences in the experimental procedure and systematic bias in the proliferation measurements. These are known problems for similar pharmacogenetic studies^47,48^ and apparently limit the achievable correlation between the predictions of the parameterized model and the Friedman data. The predicted proliferation from the mechanistic model achieves comparable agreement with the Friedman data for both individual drug treatments (r=0.35) and combinatorial drug treatments (r=0.26) (Fig. 6). Accordingly, for the combinatorial drug treatments, the model achieved a correlation with the Friedman data only 20% lower than the agreement between the datasets (0.26/0.33=0.79 compared to 1). The correlation difference between single and combination treatment was not statistically significant. We conclude that the prediction accuracy for combinatorial treatments of the mechanistic model is reasonable, given the reproducibility of the proliferation data.

**Figure 6.**
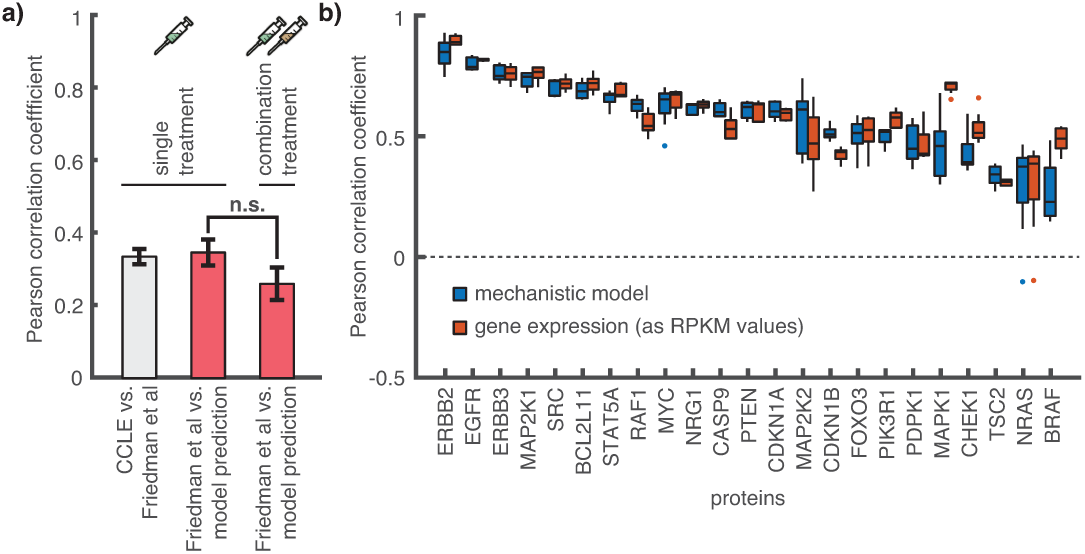
Assessment of model prediction for combination treatment dataset. (a) Correlation coefficients of the mechanistic model and data interpolation on the single and drug combination measurements by Friedman et al.^46^ evaluated for cell lines contained in the training sets of the cross-validation. Error bars show the standard error. (b) Pearson correlation coefficients of the mechanistic model and gene expression levels (as reads per kilobase of transcript per Million mapped reads (RPKM)) values with proteomic data from the MD Anderson Cell Line Project^49^ (For details see Online Methods, Section Validation). For the mechanistic model the correlation was evaluated for cell lines contained in the training sets of the cross-validation.

### Protein abundances predicted using mechanistic model

For the estimation of the model parameters only proliferation measurements were employed. As expected, our assessment of model uncertainties revealed that this limits the prediction accuracy (Online Methods, Section Uncertainty Analysis). The analysis suggested a low reliability of parameter estimates (Supplementary Fig. 2a), a higher reliability of prediction for protein abundances (Supplementary Fig. 2b) and the highest reliability for the proliferation readout (Supplementary Fig. 2c). To determine the accuracy of the predicted protein abundances in the untreated condition, we compared them to the measurements contained in the MD Anderson Cell Line Project (MCLP)^49^. The MCLP provides normalized Reverse Phase Protein Array (RPPA) data for 33 proteins described by the mechanistic model in 68 of the cell lines considered for the parameterization. We implemented the same normalization for the simulation and considered all proteins that were measured in at least 10 of the considered cell lines. The mechanistic model achieved an average correlation of r=0.57±0.03 (Figure 6b). This is similar to the correlation between gene expression and RPPA data (Figure 6b). We conclude that the individualization of the model with cell-specific gene expression levels allows for a reasonable prediction of the protein levels, despite the dependence on a large number of kinetic parameters, such as degradation rates, that were not constrained by any molecular data.

### Discussion

We generated a large-scale mechanistic model that integrates large amounts of prior knowledge and expands upon previous large-scale models^16,18^ by implementing kinetic rate laws, mutation variants of key regulators and possibilities for individualization. As the model can be individualized to particular cell lines and covers many relevant driver mutations, the model provides a valuable resource for analysis of various cancer types and drug treatments.

To parameterize this model, we established a computational framework that provides scalability with respect to the number of parameters and number of state variables, and employs parallelization to handle the large number of experimental conditions. The final wall time requirement for all optimization runs (∼4.10^3^ hours) was more than one order of magnitude lower than the wall time required for a single gradient evaluation using established methods (∼6.10^4^ hours). This allowed, to the best of our knowledge for the first time, the parameterization of a large-scale mechanistic model from experimental data from over 100 cell lines, each under dozens of experimental conditions. The computational efficiency of the approach renders iterative rounds of optimization, hypothesis generation and model refinement of large-scale mechanistic models and multiple (heterogeneous datasets) feasible in a reasonable time frame. Our implementation of the methods is available as Supplementary File 1 and can be freely reused by other research groups.

The assessment of the parameterized model revealed that the prediction of cell proliferation – a key readout to cancer therapy – is accurate for single drug treatments. Hence, the large-scale mechanistic model we derived and parameterized can predict the drug response of cancer cell lines from sequencing data. This is in our opinion a result of combining extensive integration of prior knowledge on network structure and reaction kinetics parameterized with our scalable methods. We illustrated the broad capabilities of the mechanistic model by predicting protein abundances and the outcome of combination treatments from single treatment proliferation measurements, neither of which is possible with statistical models.

Our analysis, however, also revealed limitations of the available datasets and the parameter estimates. Firstly, for combination treatments, the weak correlation of the available datasets^37^,46 limited the validation of our model and highlighted the need for accurate, reproducible phenotypic characterizations. Secondly, the parameterization of the model using only proliferation data resulted in large parameter uncertainty, which suggests that inclusion of proteomic and phosphoproteomic data in the training process will be necessary to render reliable predictions on the molecular level feasible. Thirdly, even more cell lines are necessary to capture the effect of driver mutations with low recurrence. However, as the model can be individualized to arbitrary cell lines and other experimental systems, it is particularly suited for the study of rare mutation patterns.

The developed model is currently limited to cell lines but can be extended in several ways. The modeling of additional intracellular processes, cell-cell communication, cancer heterogeneity or pharmacokinetics might improve the prediction of patients’ response. To obtain a refined description, our model could be integrated with agent-based models for tumor growth^28,50^ or physiology-based pharmacokinetic models^51^. The resulting models could provide valuable tools for the identification of novel drug targets^3^, virtual clinical trials^52^ and personalized medicine. The extensive mechanistic modeling of biological processes will therefore be an important area of future research.

## Acknowledgements

This work was supported by the German Research Foundation (DFG) through the Graduate School of Quantitative Biosciences Munich (QBM; F.F.), the European Union’s Horizon 2020 research and innovation program (CanPathPro; Grant No. 686282; A.M., B.L., C.W., D.W., J.H., L.S. and M.S.), the German Federal Ministry of Education and Research (BMBF) within the SYS-Stomach project (Grant No. 01ZX1310B; J.H.) and the Postdoctoral Fellowship Program of the Helmholtz Zentrum München (J.H.).

## Author Contributions

A.M. B.L., C.W., H.L. and T.K. designed the model. F.F. and J.H. designed the methods for numerical simulation, parameter optimization und uncertainty analysis. C.W., F.F., F.J.T., J.H., M.H. and T.K. designed the experiments. A.S., J.L., H.H., H.L., M.S. and T.K wrote and ran the code for model/data mapping and integrated and assembled model input data. D.W., F.F., J.H. and L.S. wrote and ran the code for the parameterization and assessment of the mechanistic model. M.H. wrote and ran the code for the parameterization and assessment of the statistical models. D.W., F.F., J.H., L.S. and M.H. analyzed output data. C.W., D.W., F.F., J.H., M.H. and T.K. wrote the manuscript. All authors discussed the results and implications and commented on the manuscript at all stages.

---

**Supplementary Figure 1.**
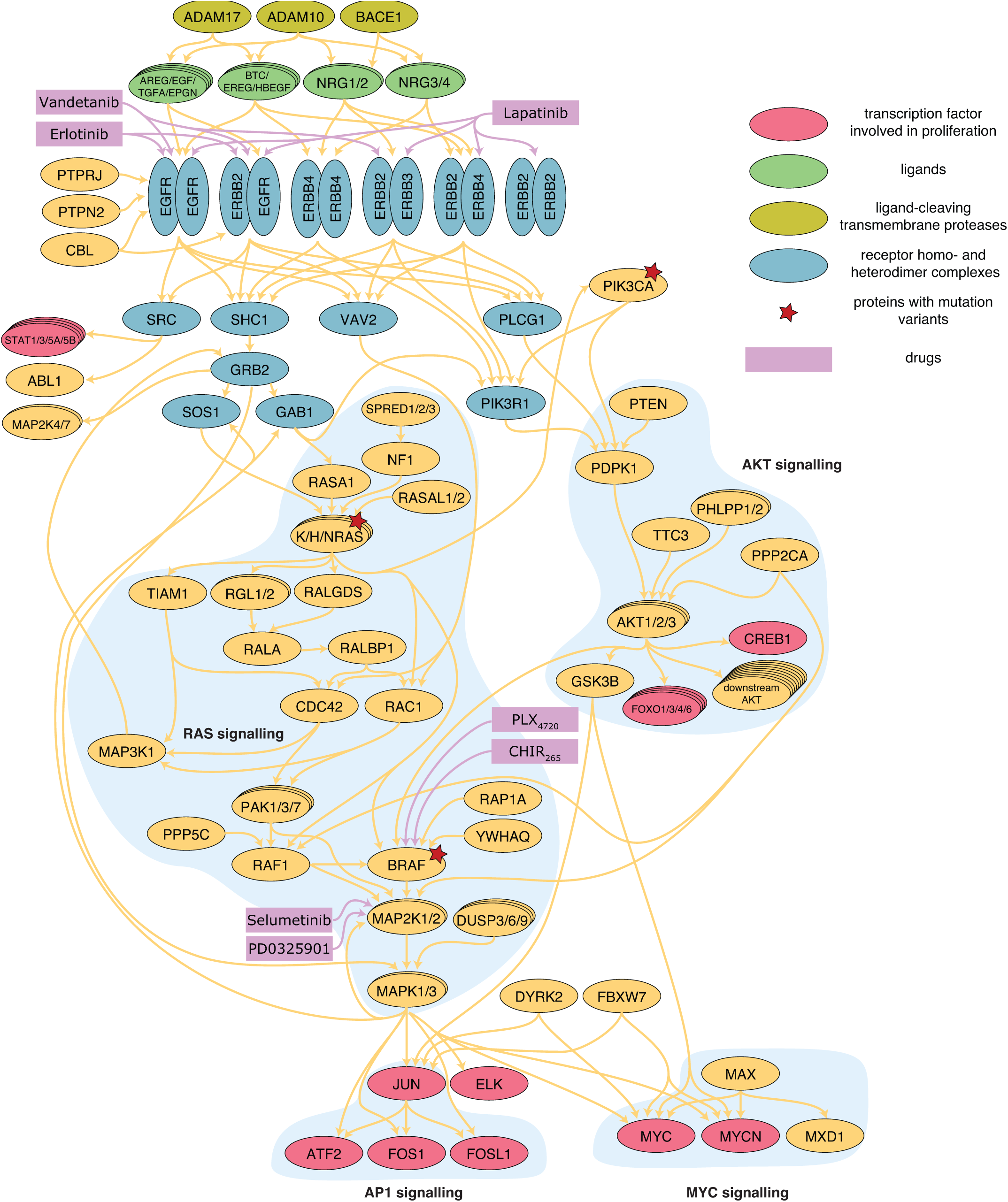
Simplified overview of the model. The figure illustrates modeled interactions. Complex formation and phosphorylation as well as activation and repression are not discriminated here. Synthesis, translocation and degradation are omitted. All species are colored according to their function.

**Supplementary Figure 2.**
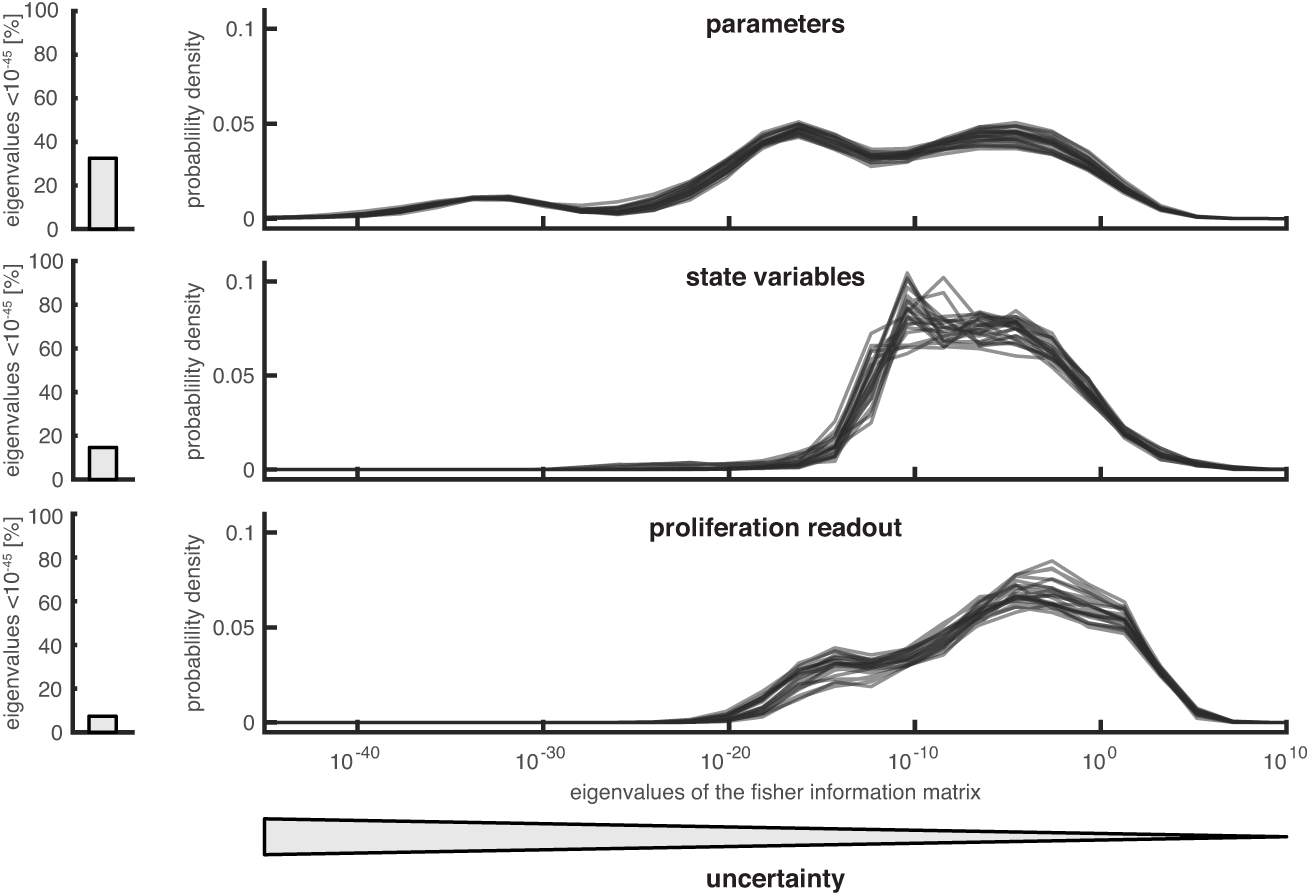
Eigenvalues densities of the Fisher Information Matrix for parameters, state variables and proliferation readouts. Small eigenvalues correspond to large uncertainties of readout combinations defined by respective eigenvectors. One line for each of the best 5 optimization runs for each of the 5 cross validation is shown. All eigenvalues below 10^-45^ are not shown in the density plot but the corresponding fraction of eigenvalues is indicated in the barplot on the left.

**Supplementary Figure 3.**
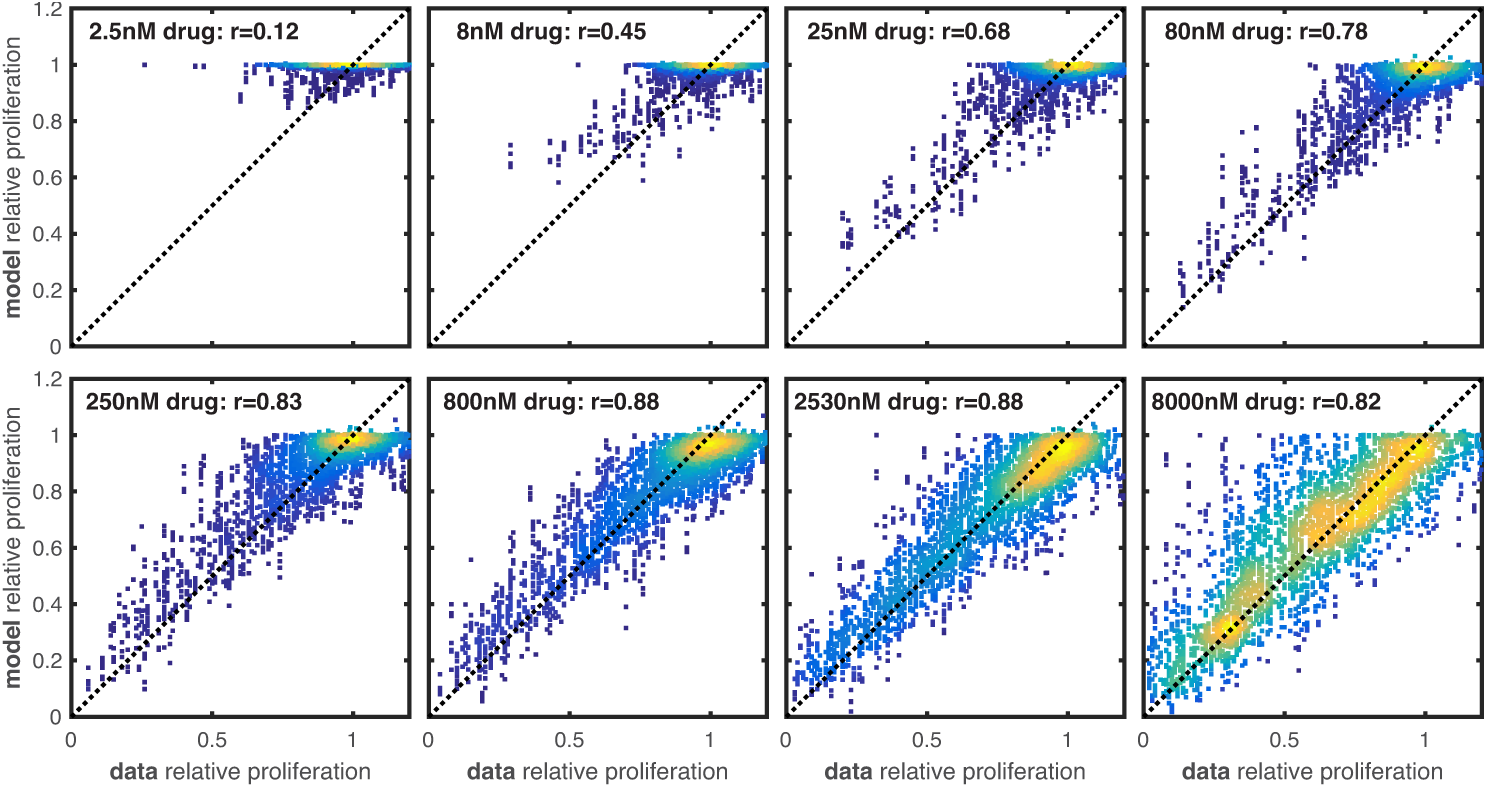
Correlation of model simulation and experimental data at all measured drug concentrations. The correlation for the drug concentrations from 250nM to 2530nM is higher than at 8000nM, which is likely due to an inflation of cell lines not responding to drugs (relative proliferation=1). For lower drug concentrations the correlation is lower than at 8000nM, which is due to the lower dynamic range of simulated and experimentally observed relative proliferation values.

**Supplementary Figure 4.**
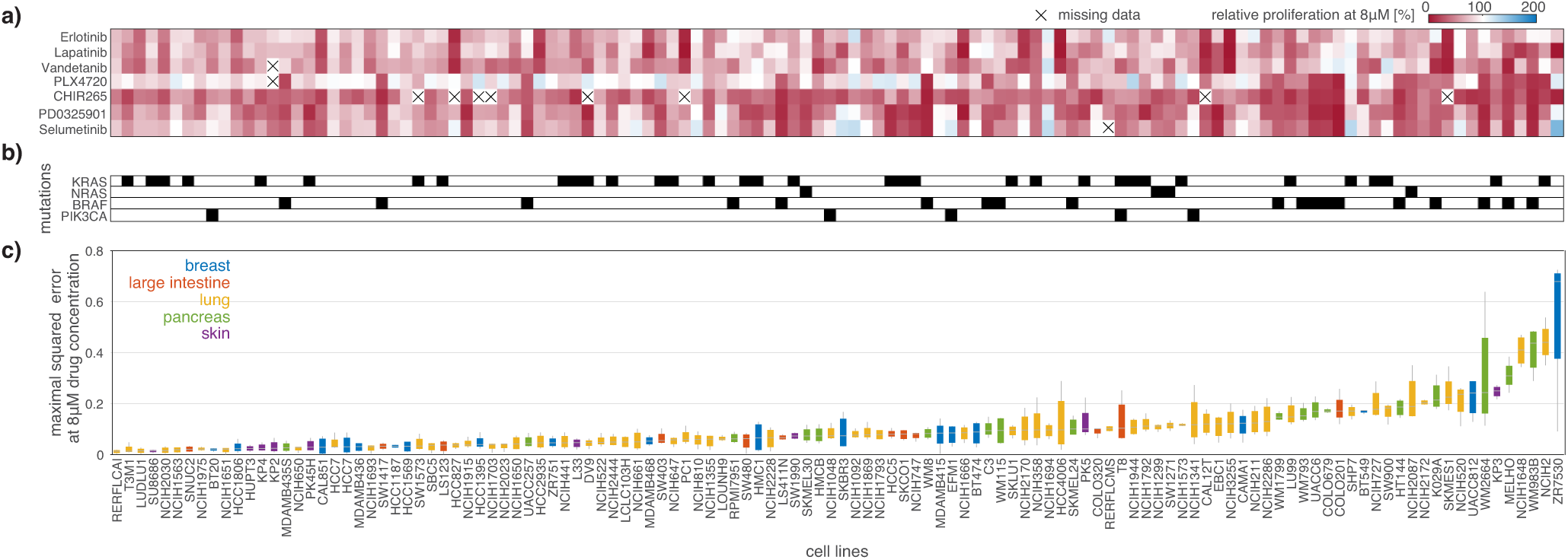
Overview of parameterization results. One column corresponds to an individual cell line. The cell lines are sorted according to the median over cross-validations of the maximal squared error at 8µM drug concentration shown in c). (a) Measured relative proliferation in response to the treatment with 8µM of the different drugs. (b) Gain-of-function mutations in the individual cell lines. Mutation status is summarized per gene and does not distinguish individual variants. (c) Boxplots of the maximal squared error at 8µM drug concentration over the 5 cross-validations. The squared error is evaluated for the median of the simulation from the 5 best optimization runs. The maximum is taken over all drugs. The boxplots are colored according to the tissue of origin of the cell lines.

**Supplementary Figure 5.**
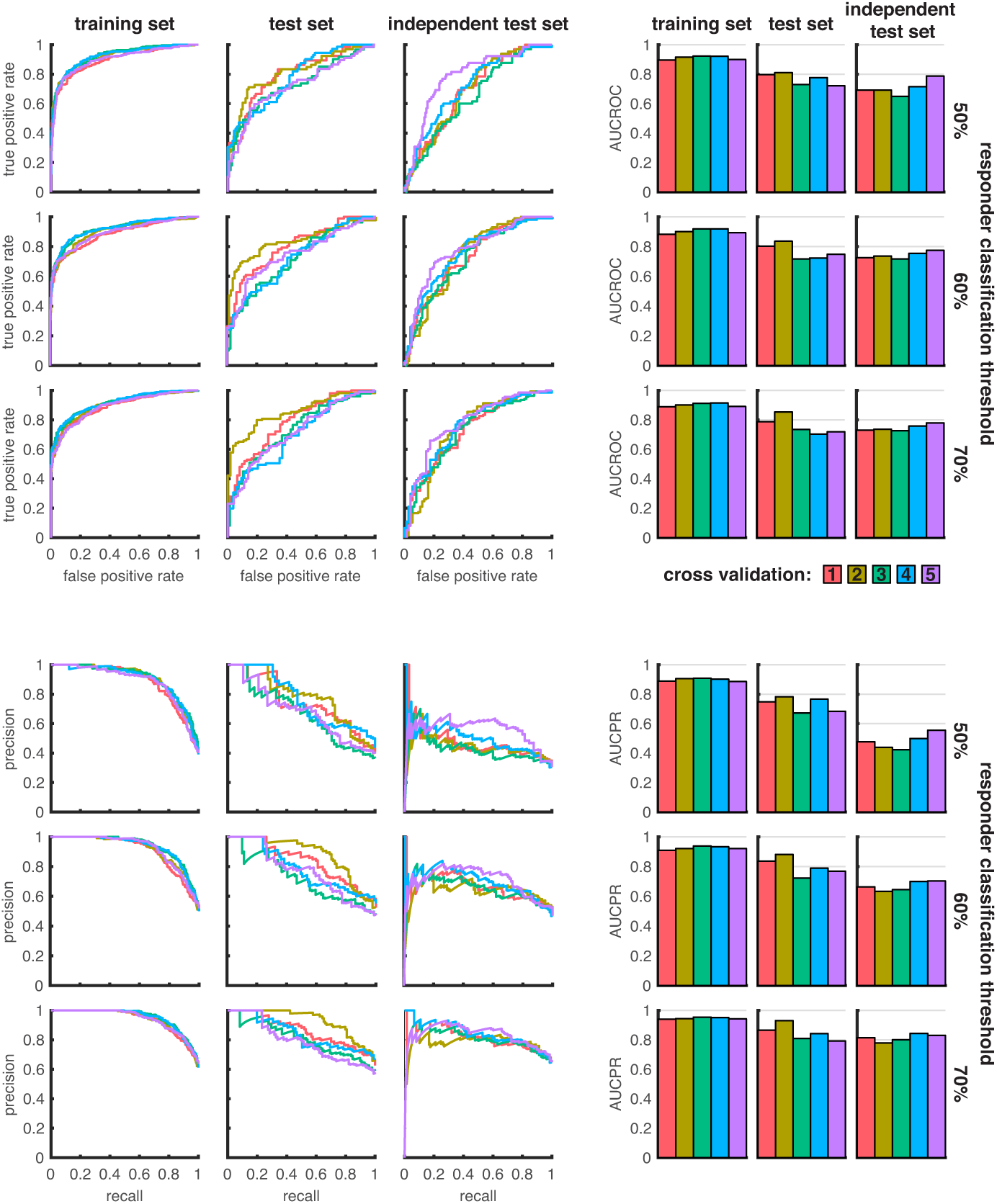
Receiver-operating-characteristic and precision-recall analysis for different datasets and classification thresholds. Every column corresponds to a dataset. Every row corresponds to a different value for the classification threshold. Colors indicate the 5 different cross-validations.

## Online Methods

### Model Development

The mechanistic model was developed using PyBioS^30^, a web-based platform for modeling of complex molecular systems. We exploited several features of PyBioS, including the modular formulation of large-scale models based on individual pathways and their interactions. For model development we employed information from ConsensusPathDB32. The information was manually curated and implemented in the model using a standard operating procedure (SOP). The SOP ensured the model quality and the compatibility of different pathway models. The model structure was refined several times by different experts to ensure highest standards. To validate the model structure a plethora of logical test was used, e.g. to certify that the known effects of growth factor stimulations are correctly captured.

The developed model is made publicly available as supplement to this manuscript. The model features an exhaustive annotation, including UniProt and Ensembl IDs. Phosphorylations are indicated in the name of the species by a preceding “P[$X;$Y;…]-“ where $X and $Y specify the phosphorylation site using a one letter amino acid code, followed by the amino acid number. Mutations are indicated in the name of the species by a preceding “MutAA[$Z]-“ where $Z specifies the mutation site using standard sequence variant nomenclature. Homodimers are indicated by a trailing “[2x]".

The SBML file encodes an ordinary differential equation (ODE) model of the form

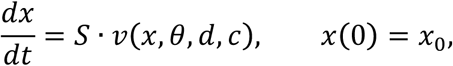

with concentration vector x and its initial condition *x*_*0*_, stoichiometric matrix *S* and flux vector *v*. The parameter vector θ provides the reaction rates, e.g. binding affinities. The vector *d* provides the drug concentrations used for simulation and the vector *c* provides the expression levels for the gene products and respective variants for a particular cell line. To consider different drug treatments and cell lines, only *d* and *c* need to be changed. The parameter vector is generic and transferable. The SBML model provides a representative parameter estimate.

In the SBML model the proliferation output variable is specified as an assignment rule. The proliferation output is computed as fraction of weighted sums of concentrations of active forms of transcription factors

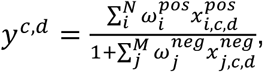

in which 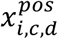 and 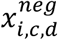 denote the concentrations of transcription factors, for a particular cell line and drug treatment combination, with a positive and negative influence on proliferation, respectively. The corresponding weights are denoted by 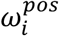 and 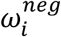 and were estimated during model parameterization. The model captures the effect of N = 12 transcription factors with positive influence: P[S63;S73]-JUN[2x], P[S252;S265]-FOSL1:P[S63;S73]-JUN, P[T69;T71]-ATF2:P[S63;S73]-JUN, P[S374;T325;T331]-FOS:P[S63;S73]-JUN P[Y701]-STAT1[2x], P[Y705]-STAT3[2x], P[Y694]-STAT5A[2x], P[Y699]-STAT5B[2x], MAX-001:P[S62]-MYCN, MAX:P[S62]-MYC, P[S324;S383]-ELK1, and P[S133]-CREB1. Furthermore, *M = 4* transcription factors with negative influence were included: FOXO1, FOXO3, FOXO4 and FOXO6. In all cases only the species with nuclear localization were considered to be active.

The model employs experimentally derived drug-target binding affinity (Kd) values for the drugs CHIR-265, erlotinib, lapatinib, PLX4720, selumetinib, sorafenib and vandetanib, which were obtained from Davis *et al.*^53^. For PD0325901 the model employs the inhibitory concentrations (IC50), which was measured in a cell-free assay by Barrett *et al.*^54^.

We note that the model includes several components that were not used in the presented analysis. This includes the option to specify gene specific scaling constants to individually adjust synthesis rates. Furthermore, the small molecular kinase inhibitor sorafenib was modeled. However, as none of the considered cell lines responded to sorafenib and as sorafenib targets several components that are not captured by the model, the corresponding response data was not considered in this study.

### CCLE Data Processing

We downloaded RNAseq BAM-files for 780 CCLE cell lines from the Cancer Genomics Hub (https://cghub.ucsc.edu/) in April 2014. The same data, including additional cell lines is now available for download in the Cancer Genomics Cloud (https://cgc.sbgenomics.com/). The gene expression values were normalized as Reads Per Kilobase of transcript per Million (RPKM) using gene models from Ensembl Release 73. Mutation data was downloaded from the CCLE data portal (https://portals.broadinstitute.org/ccle/data/, file CCLE_hybrid_capture1650_hg19_NoCommonSNPs_NoNeutralVariants_CDS_2012.05.07. maf). RNA allele frequencies for the mutations were determined from the downloaded RNAseq BAM-files using SAMtools mpileup (http://www.htslib.org). Drug response data were downloaded from https://portals.broadinstitute.org/ccle/downloadFile/DefaultSystemRoot/exp_10/ds_27/CCLE_NP24.2009_Drug_data_2015.02.24.csv?downloadff=true&fileId=20777.

Of the 780 cell lines, for which we processed RPKM values, 123 originated from the tissues breast, large-intestine, lung, pancreas and skin which were considered in the training/test data-set and 31 originated from the tissues kidney, soft tissue, ovary and stomach which were considered for the independent test set. For the training/test data we considered 120 of the 123 available cell lines to ensure equally sized training and test sets in all cross-validations.

To generate test and training datasets from the processed CCLE data, we performed 20-80% splits on the cell-line level, which yielded 5 training sets with 96 cell lines and test sets with 24 cell lines. The split was performed such that the tissue distribution in the individual training sets is maximally similar. The number of experimental condition in the training sets varies from 5390 to 5403 due to incomplete data for individual cell lines.

### Numerical Simulation and Gradient Evaluation

The compilation and numerical simulation of the model was performed using the MATLAB toolbox AMICI^40^ (http://dx.doi.org/10.5281/zenodo.579891). AMICI employs the backward differentiation method implemented in the SUNDIALS solver package^55^. We used the KLU linear solver with AMD reordering and relative and absolute error tolerance 10^-8^. As the proliferation measurement was taken after 72 to 84 hours, we assumed that the state of the model reached a steady state. To find the steady state for the untreated condition of a cell line, the forward simulation was initialized with zero and run until the regularized maximal absolute relative derivative was smaller than 10^-6^,

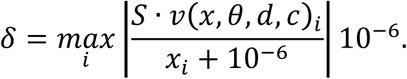

For all treated conditions of a cell line, the forward simulation was initialized with the steady state of the corresponding untreated condition.

The objective function gradient was computed using adjoint sensitivity analysis^40^. As the model only possesses a single model output, the proliferation *y*, we computed the sensitivity of this output. From this output sensitivity, we computed the objective function gradient and the Fisher Information Matrix (FIM). The FIM is not accessible when adjoint sensitivity analysis is used to directly compute the objective function gradient.

The forward and backward simulation of experimental conditions was parallelized using the MATLAB command *parfor*, which implements OpenMP parallelization. As our cluster infrastructure features 8 core nodes, we parallelized each gradient computation over 8 cores (1 master, 7 workers), thereby avoiding inter-node communication overhead. The different local optimizations were performed on different nodes.

### Numerical Benchmark

To compare different methods for gradient evaluation, we assessed the computation time for a single gradient evaluation on the full training set. For sequential and parallel gradient evaluation using adjoint sensitivity analysis, we measured the computation time. As this would have been too time consuming for forward sensitivities, we first assured that the computation time for individual experimental conditions is comparable and then extrapolated to all experimental conditions. The computation time was evaluated on the training set of the first cross-validation for 10 randomly sampled parameter vectors. For the difference between forward and adjoint sensitivity analysis and sparse and dense solvers, we only evaluated the simulation time for the untreated condition of a single cell-line. The performance was evaluated based on 100 samples with a randomly drawn parameter vector and a randomly drawn cell-line. The computation time was then normalized such that the median for the sparse adjoint approach matched the computation time for the full training set.

### Parameterization

To estimate the model parameters θ, we used the measurement data for the proliferation in the treated condition relative to the untreated condition, 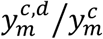, provided in the CCLE dataset. These data were fitted using a sum-of-squared-residuals objective function

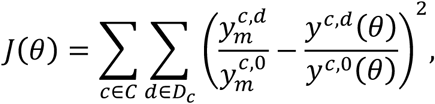

in which *c ∈ C* is cell-line specific and *d ∈ D*_C_ denotes the drug treatment. This objective function is equivalent to the negative log-likelihood function under the assumption of additive independent and identically distributed standard normally distributed measurement noise. To minimize the objective function we used multi-start local optimization^26^ implemented in the MATLAB toolbox PESTO (http://dx.doi.org/10.5281/zenodo.579890). Parameters were constrained to a [10^-2^,10^2^] hypercube. For each local optimization run, parameters were drawn from this hypercube, followed by 100 optimization iterations of the MATLAB fmincon interior-point algorithm. The gradient of the objective function was computed from adjoint sensitivities and supplied to the interior-point algorithm. A high-performance-computing-ready standalone executable was generated from the parameterization pipeline implemented in MATLAB using the MATLAB Compiler toolbox. For every cross-validation we performed 10 local optimization runs. As no communication was necessary between optimization runs, each could be submitted as a separate job to the cluster. In total we submitted 50 jobs using 8 cores each, resulting in a total parallelization over 400 cores.

### Ensemble Averaging

We used ensemble averaging to reduce the effect of overfitting and the variance of predictors. For the mechanistic model we used an ensemble model based on five optimization runs that achieved the lowest objective function value. The parameter values from these optimization runs were then used to simulate the model individualized to cell lines from the test and independent test set. For quantitative predictions the median of the five simulations was used and for classification a majority vote was used. The ensemble averaging was solely based on results from the training set. The test and independent test set were only used for validation.

### Uncertainty Analysis

We assessed the uncertainty of parameters using the eigenvalue spectrum of the Fisher Information Matrix (FIM). Small eigenvalues indicate large uncertainties in the direction of the respective eigenvector while large eigenvalues indicate small uncertainties. The eigenvalue spectrum was evaluated for the best 5 optimization runs for every cross-validation.

The FIM was computed by summing the dyadic product of adjoint sensitivities over all experimental conditions

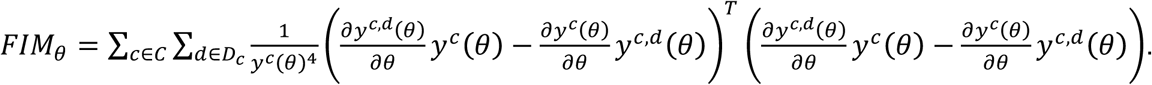

As the number of experimental conditions (∼5.400) exceeds the number of parameters (∼4,100) the FIM could theoretically have full rank.

For parameter derived readouts *z*, such as proliferation readouts as well as state variables, a similar quantification of the uncertainty is possible by considering a transformation *FIM*_*z*_ of the FIM. The transformation is obtained by multiplication with the respective parameter derivatives

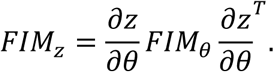

For state variables the formula for steady state parameter derivatives were computed according to the implicit function theorem

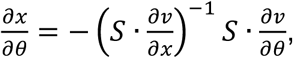

assuming that the system is in steady state,

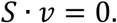

For the *FIM*_*z*_ for state variables, we only considered state variables with non-zero simulated steady state. The state variables with steady state equal to zero correspond to molecular species that are not expressed.

### Statistical Methods

For the comparison of model performances, we trained a series of statistical models for the prediction of response to treatment, based on the exact same training data sets, cross-validation setup and test data sets that were used for the mechanistic model. Responder and non-responder cell lines were defined for each drug by applying the threshold 0.5 on the proliferation at the highest dose used for the drug (in addition results for thresholds 0.7 and 0.9 were also obtained). The classification of cell lines into responders and non-responders was evaluated for three sets of input variables: 1) mutation genotype data; 2) gene expression data; and 3) genotype and gene expression data. In addition we also provided the network topology as input to some of the classifiers. Model training was performed by nested cross-validation, where the outer 5-fold cross validation loop split the data into training set (80% of the data) and test set (20% of the data). For each classifier and each training set we estimated the model parameters by optimizing the classification performance in the inner cross validation loop splitting the training data again into training and validation sets with balanced class labels or by bootstrapping of the training data.

We used the R implementations of the following classifiers: 1) logistic regression with LASSO penalty (glmnet package); 2) Random Forest (randomForest package); 3) graph regularized logistic regression (glmgraph package); and 4) logistic regression with LASSO penalty on augmented data, including additional interaction terms. The interaction terms were defined based on the network topology used in the mechanistic model. The adjacency matrix was extracted from the Jacobian of the right hand side of the differential equation. For a pair of genes the Jacobian was reduced to rows or columns corresponding species that include the corresponding proteins in any form (phosphorylated, cleaved or bound). Two genes were defined to be adjacent when the corresponding submatrix of the Jacobian has at least one non-zero entry. We augmented the data set either with all pairwise interactions (product) between variables of the same type (genotype or gene expression) or interactions between genes that are connected by paths in the network not longer than 1, 2 or 3 steps. The optimal parameter *λ* for LASSO models, *λ*_1_and *λ*_2_ for the graph regularized LASSO model were selected as the largest *λ* that produces an AUC ROC within one standard error of the maximum AUC ROC^56^ in a 8-fold inner cross validation. Random Forest classifiers were trained by selecting parameters that minimize the out of bag error^43^. We optimized over the number of variables randomly sampled as candidates for each split, the number of trees in the forest ranging from 50 to 500 and the maximum leaf node size criterion ranging from 1 to 1/3 of the data set. All classifiers were then applied to the test set and performance was assessed as the area under the ROC curve.

Finally, we used the classifiers trained in each round of the cross validation and applied them to an independent test set. Performance was assessed as the area under the ROC curve and averaged over the five cross validation rounds.

### Validation

#### Area Under the Curve

For the quantification of the classification accuracy for the mechanistic model we computed receiver-operating-characteristic (ROC) curves and precision-recall (PR) curves. For the data and model, a cell line was classified as responder to a drug when the measured/simulated relative proliferation at 8µM was smaller or equal to the specified threshold. For the data, this threshold was fixed to 0.5 and for the model the threshold was continuously varied between 0 and 1 and the precision, sensitivity and specificity was evaluated for every threshold value. From these evaluations the ROC and PR curves were constructed and the area under the curve was evaluated using a trapezoidal rule.

Exactly the same split in training and test set was applied for all statistical models and the mechanistic model. For both approaches the test and independent test sets were never used for parameterization. For the statistical model the AUC was averaged over drugs and cross-validation as individual models were constructed for every drug. For the mechanistic model the AUC was averaged only over cross-validations as a single model for all drugs could be constructed.

#### Combination Therapy

The data by Friedman *et al.*^46^ includes measurements for single and paired treatments at low and high concentrations for the drugs selumetinib, CHIR-265, erlotinib, lapatinib and PLX4720. For the paired treatment, the experiment was repeated twice and the average value was used for our analysis. As the treatment concentrations employed in the two studies did not agree, we interpolated measurements from the CCLE data to concentrations employed in the Friedman dataset. The interpolation was performed in logarithmic concentration space but results were comparable for linear concentration space. For every cross validation we only considered the measurements from cell lines which were also contained in the training set.

#### Proteomics Measurements

For the comparison of model prediction and RPPA data from the MD Anderson Cell Lines Project, we computed the median total protein concentrations for the 5 best optimization runs. To compute total protein concentrations, we computed the sum of concentrations of all protein and complex species that contain a specific protein, taking into account the individual stoichiometry. As the RPPA data in the MD Anderson Cell Lines Project are normalized, we also normalized the simulation data using the same procedure: 1) The simulated concentrations were log_2_ transformed; 2) The median concentration over all cell lines was subtracted for every protein and then the median of all concentrations was subtracted for every cell-line. While this procedure resembles the procedure described in the manuscript providing the data, we note that the model only captures a fraction of the proteins and the normalization might therefore be suboptimal. For the RPKM data, only a log_2_ transformation was applied. The correlation coefficients of normalized data and predictions were computed for all proteins that were measured in at least 10 cell lines. For every cross validation we only considered the measurements from cell lines which were also contained in the training set.

The considered dataset only provides measurements for the untreated condition. However, the parameterization was performed based on relative proliferation values, which is computed based on the ratio of sums of active transcription factor concentrations in the model. Accordingly, the training data provides some information about the ratio of concentrations in treated and untreated condition, but little information about absolute concentrations in the treated and untreated condition.

